# Sex- and age- dependent effect of pre-gestational chronic stress and mirtazapine treatment on neurobehavioral development of offspring

**DOI:** 10.1101/2021.08.04.455108

**Authors:** Viñas-Noguera Mireia, Csatlósová Kristína, Šimončičová Eva, Bögi Ezster, Ujházy Eduard, Dubovický Michal, Belovičová Kristína

## Abstract

Hormonal fluctuations, such as the perinatal period, may increase susceptibility of women to depression, which in turn exert a negative impact on child’s neurodevelopment, becoming a risk factor in development of neuropsychiatric disorders. Moreover, the use of antidepressants during this critical period presents a serious health concern for both the mother and the child, due to the consequences of treatment in terms of the reliability and safety for the proper neurodevelopment of the organism being not well known. Atypical antidepressants, such as mirtazapine, that targets both serotonergic and noradrenergic systems in the central nervous system (CNS), represent a novel focus of research due to its unique pharmacological profile. The aim of this work was to study the effects of maternal depression and/or perinatal antidepressant mirtazapine treatment on the neurobehavioral development of the offspring. Pre-gestationally chronically stressed or non-stressed Wistar rat dams were treated with either mirtazapine (10 mg/kg/day) or vehicle during pregnancy and lactation followed by analysis of offspring’s behavior at juvenile and adolescent age. We found mirtazapine induced alterations of nursing behavior. In offspring, pregestational stress (PS) had an anxiogenic effect on adolescent males and increased their active behavior in forced swim test. Interaction between pregestational stress and mirtazapine treatment variously induced anxiolytic changes of juvenile and adolescent females and impairment of spatial memory in adolescent females as well. Hippocampal density of synaptophysin, pre-synaptic protein marker, was decreased mainly by mirtazapine treatment. In conclusion, our results show mirtazapine induced alterations in maternal behavior and several sex- and age-dependent changes in neurobehavioral development of offspring caused by both prenatal mirtazapine treatment and/or chronic pregestational stress.

## INTRODUCTION

An estimation of 17% men and 25% women experience an episode of major depressive disorder (MDD) at least once in their life (1). Higher susceptibility of women to depression may arise from the increased vulnerability caused by periods of hormonal fluctuation, such as the perinatal period (2,3). Perinatal depression has been reported to exert a negative impact in children neurodevelopment, becoming a risk factor in developing neuropsychiatric disorders (4–6).

Second generation antidepressants (SGA), such as selective serotonin reuptake inhibitors (SSRIs) (fluoxetine, sertraline) or serotonin-norepinephrine reuptake inhibitors (SNRIs) (venlafaxine) introduced in Europe in the 1980s, are recorded as the first line antidepressant treatment during pregnancy by most medication guides (7,8). The World Health Organization (WHO) estimates that depression is a leading cause of disability worldwide (9). However, the main concern for specialists is the impact of the treatment on the developing fetus/child, namely risk of potential congenital malformations, neonatal withdrawal symptoms or poor neonatal adaptation syndrome as well as long-term neurodevelopmental consequences (10–13). In addition, the delay in treatment efficacy and the presence of many side effects lead researchers to investigate alterative antidepressants with better efficacy, faster onset of action and lesser counterproductive reactions.

Mirtazapine (MIR) (Fig 1) has a tetra-cyclic chemical structure with molecular weight of 265.36 and belongs to the piperazino-azepine group of compounds. It is a new generation antidepressant that has a different mechanism of action than SGAs, targeting both the serotonergic and noradrenergic systems in the CNS. It is a noradrenaline (NE) and specific serotonin (5-HT) antidepressant (NaSSA) that acts as an antagonist at central α2-adrenergic inhibitory autoreceptors and heteroreceptors, as well as at the 5-HT_2_ and 5-HT_3_ receptors. MIR enhances the release and availability of NE by blocking presynaptic inhibitory α2-autoreceptors and enhances the 5-HT release by antagonism of the α2-heteroreceptors in the serotonergic nerve terminals and simultaneously blockade of postsynaptic 5-HT_2_ and 5-HT_3_ receptors. Thanks to this double activity, MIR is suggested to induce earlier onset of antidepressant effects avoiding the serotonergic related side effects such as high body temperature, agitation, increased reflexes, tremor, sweating, dilated pupils, and diarrhea (14). However, it may induce an enhanced body weight gain and sleepiness (15,16). Mirtazapine has low *in vitro* affinity for central and peripheral dopaminergic, cholinergic, and muscarinic receptors, but high affinity for central and peripheral histamine H1 receptors. However, it appears that the antihistaminergic effects of the drug are counteracted by noradrenergic transmission when the drug is commenced at dosages ≥15 mg/day, i.e. within the recommended dosage range (17). Initial dose of MIR is 15 mg/day orally once a day at bedtime and maintenance dose represents 15 to 45 mg orally once a day (18). Even though MIR seems to represent a plausible alternative of antidepressant medication during gestation and there is no evidence that use of mirtazapine in pregnancy causes birth defects, preterm birth, or low infant birth weight. While the evidence for other pregnancy outcomes is also reassuring, only small numbers of women have been studied and number of animal studies is not sufficient (19).

**Figure 1.**
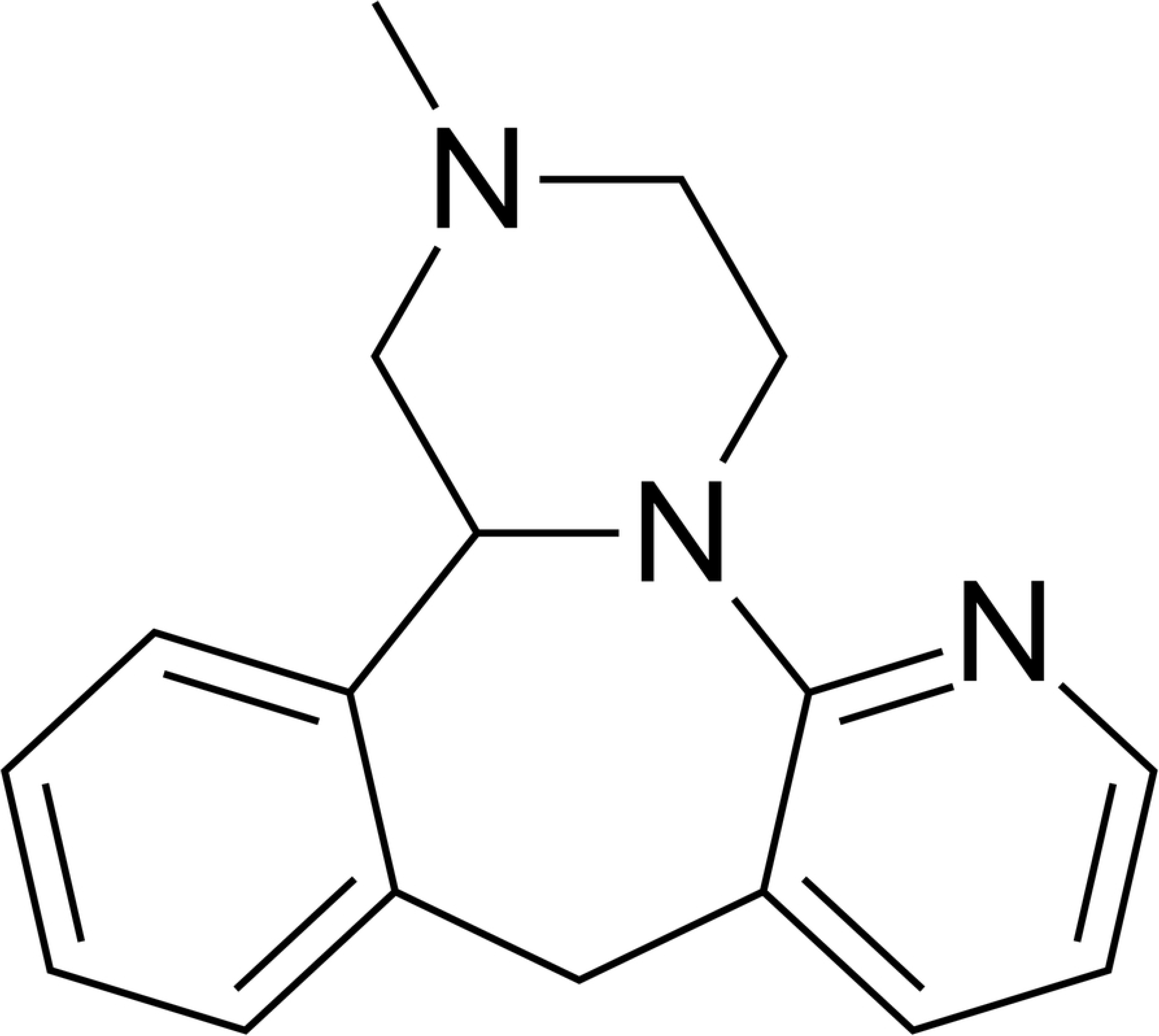
Chemical structure of mirtazapine.

Despite lacking efficient translatability, several animal models, including chronic unpredictable stress, have been investigated for decades to evaluate depression-like behavioral changes and their potential mechanisms of action. Studies in rodents show that chronic pre-gestational and prenatal maternal stress, which disrupt the maternal endocrine, nervous and immune systems, can induce long-term alterations in the synaptic structure and so impact the behavioral outcomes in the offspring (6,20). Prior to regulatory roles of serotonin, norepinephrine and dopamine neurotransmitters in adult brain, monoamines play an important role in the fetal maturation of the brain, such as - during neuronal proliferation, migration and differentiation, myelinization and synaptogenesis (21). The exposure to stressful situations early in life may disable the optimal structural and functional development of hippocampus, due to the damaging action of excessive corticosterone concentration and dysregulation of monoamines (11,22). Persisting high levels of stress are thought to result in loss of synapses in circuits underlying affective and cognitive processes. These reductions are presumed to contribute to the symptoms of depression associated with major depressive disorder (23). Synaptophysin is a synaptic vesicle glycoprotein, which immunoreactivity is present in a punctate pattern in the hippocampus and has been used as a presynaptic marker for quantification of synapses (24). The hippocampal formation consists of several histologically distinguishable modules, such as Cornu Ammonis (CA) regions (CA1, CA2, CA3, CA4), dentate gyrus (DG), presubiculum, and subiculum. These regions of the hippocampus are associated with different functions (e.g. memory encoding and retrieval) and may be specifically disrupted in various diseases (25). Granule cells, the main output cells of the DG, send axons (mossy fibers) through CA4 to CA3 and innervate a small number of pyramidal cells and a disproportionally large number of interneurons CA3 pyramidal cells then form recurrent excitatory network and send axons to CA1. Theoretical work, as well as anatomical, physiological, and behavioral experiments support the idea that the DG-CA3 system performs the pattern separation and the pattern completion of the inputs to the hippocampus, operations needed for memory encoding and retrieval (26). Purpose of the hippocampus happens to be severely affected by early life stress, predisposing the individual to an impaired reactivity when exposed to adverse environmental stimuli (27). In our previous study, we have shown that pre-gestational maternal stress may affect hippocampus at the time of birth by increasing the resting membrane potential, suppressing depolarization-activated action potential firing, and increasing the spontaneous activity of hippocampal cells from newborn rat offspring (28), but pre-gestational stress induced changes in different subregions of hippocampus are not thoroughly known. However, antidepressant treatment may facilitate the challenged neurogenesis by, upregulating the hippocampal concentration of glucocorticoid receptors, which help to attenuate the hyperactivity of the HPA axis and inducing morphological changes in the neuronal network (11,27,29). Some studies suggest that developmental fluoxetine (SSRI) exposure of prenatally stressed male and female offspring reversed the effect of stress on the number of immature neurons in the dentate gyrus (DG), with effects being more prevalent in adult male offspring (30,31). However, knowledge of mirtazapine treatment’s impact on pregnancy and lactation hippocampal neurogenesis of juvenile, adolescent and adult offspring during last days of pregnancy and for following 2 weeks postpartum (PP), is very limited.

The aim of the present study was to determine the possible implications that pre-gestational chronic stress have on the behavioral and neurodevelopmental outputs in the offspring of both sexes during juvenile and adolescent age and to investigate the consequences of administration of the new generation antidepressant mirtazapine on these variables aiming our focus on functional brain developments that starts day 10 PP (32–35).

## MATERIALS AND METHODS

### Animal breeding

Female nulliparous Wistar rats (initial weight 200-220 g, age 2-3 months, n=44) used in this study were obtained from the Department of Toxicology and Laboratory Animal Breeding Station of the Institute of Experimental Pharmacology and Toxicology, Centre of Experimental Medicine of the Slovak Academy of Sciences, Dobra Voda, Slovak Republic. After 7 days of acclimatization, females were randomly assigned to stress or non-stress groups. Animals in the stress group were exposed to unpredictable stressors of mild intensity for a total of 3 weeks. One week after the end of the stress procedure, females were mated with males in the ratio 3:1. The presence of spermatozoa in vaginal smears was considered day 0 of gestation. On day 15 of gestation, the females were separated and housed individually. The animals had *ad libitum* access to food pellets and water and were kept in a temperature and humidity-controlled room (20–24°C and relative humidity 50–60%) with 12/12 hours of light/dark cycle. The experiments were conducted in compliance with the Principles of Laboratory Animal Care issued by the Ethical Committee of the Institute of Experimental Pharmacology and Toxicology, Slovak Academy of Sciences, Bratislava, and the experimental design was approved by the State Veterinary and Food Administration of the Slovak Republic.

One day after birth, litters were culled to four males and four females (with eight offspring per cage). Reproductive variables were recorded right after the birth. The offspring were weaned on post-partum day 21 and housed in litter groups of same-sex four animals per cage. No more than two pups from the same mother per group (n= 6-8 animals/group) were used for behavioral testing.

### Chronic unpredictable stress schedule

In order to not expose females to stressors during pregnancy but rather study ongoing effects of chronic stress, females were assigned to stress or non-stress groups prior to breeding (Fig 2). The animals were exposed to 1-2 stressors per day. The list of stressors included: 1) Overcrowding - 6 rats housed together (24h); 2) Exposure to damp bedding (12h); 3) Food deprivation (24h); 4) Predator stress – cloth with cat odor (10h); 5) Water deprivation (12h); 6) Cage decline at 45-degree angle (6h); 7) Strobe light- flicker lights (12h).

**Figure 2.**
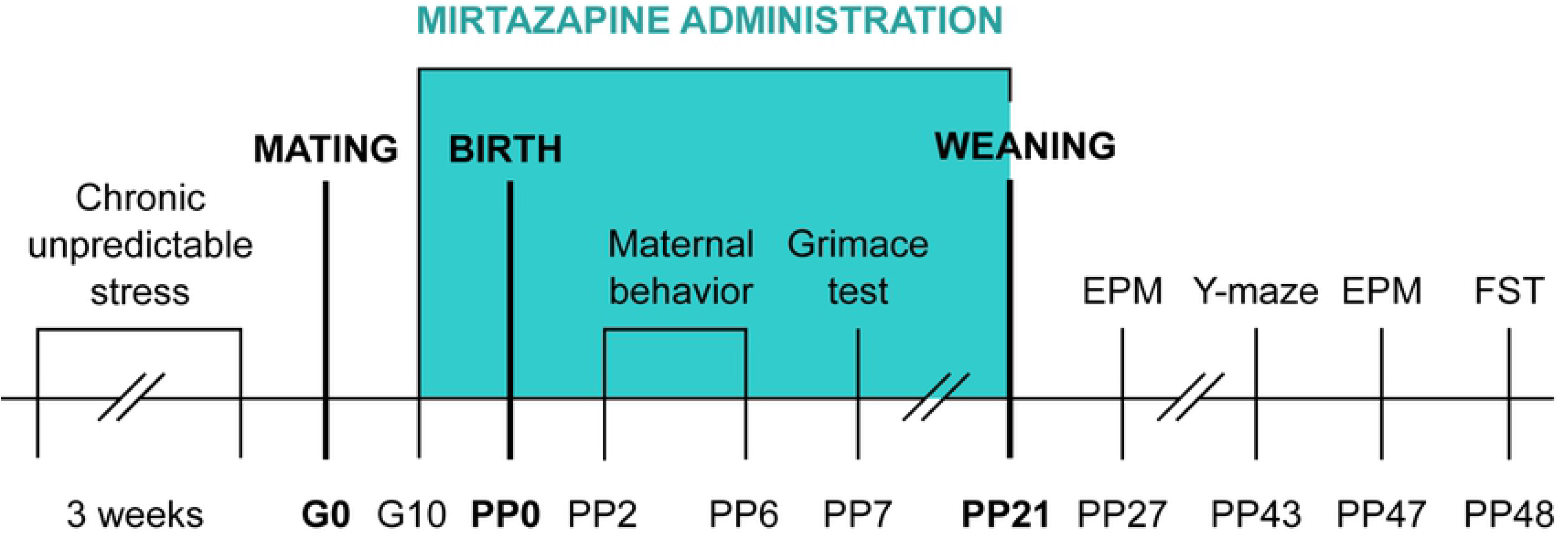
Schedule of the experiment. G-gestation day; PP-post-partum day; EPM-elevated plus maze; FST-forced swim test.

### Mirtazapine treatment

Mirtazapine (Mikrochem Trade spol. s r.o.) with a molecular weight of 265.36 was diluted in citric acid and administered orally via the wafer biscuit (size of 1 cm^3^) to pregnant rats from day 10 of gestation until sacrifice at the clinically relevant dose of 10 mg/kg once per day. Pups were receiving mirtazapine via mother until weaning on day 21 *post-partum* (PP). This dose was chosen based on a body surface area normalization (BSA) conversion used to determine the starting human dose extrapolated from animal studies. The Km factor, which is the ratio of body weight (kg) to body surface area (m^2^), is used to convert doses expressed in mg/kg to units of mg/m^2^ (36). Dose 10 mg/kg a day was used to simulate low dose end in humans. The dams from control groups received 1 cm^3^ of wafer biscuit once a day filled with vehicle (water). Feeding was completed under investigator’s supervision to ensure the dam consumed the entire biscuit. Pregnant dams were randomly divided into 4 groups: non-stress + VEHIC (vehicle), non-stress + MIR, stress + VEHIC, stress + MIR.

### Rat Grimace Scale

Mothers were placed in individual cages for 10 min and each minute 4 pictures were taken from each rat from front with a Nixon professional camera. Animals were tested 5 weeks after CUS. Each picture was evaluated by a score from 0 to 2 according to the following four action units (37):

1. Orbital tightening: rats in pain display narrowing of the orbital area which manifests as partial or complete eye closure or squeezing.
2. Nose/cheek flattening: less bulging of nose and cheek and absence of crease between cheek and whisker pads.
3. Ear changes: ears in pain tend to fold, curl and angle forwards or outwards, and the space between ears is wider.
4. Whisker change: whiskers move forward (away from face) and tend to bunch.

### Behavioral tests

#### Maternal behavior

Mothers were observed for 5 consecutive days from PD2 until PD6 two times per day for 5 min based on previous literature (20). Observations took place in the morning (between 8:30 a.m. and 10.00 a.m.) and the afternoon (between 2:30 p.m. and 3:30 p.m.). Time and number of bouts in the following maternal behaviors were recorded: licking; licking/nursing; nursing (arched-back nursing, blanket nursing, and/or passive nursing); nest building and time off the pups. Differences between groups in maternal behaviors were assessed using total time spent in these maternal behaviors.

#### Forced swim test (FST)

The FST consists of a container filled with water where the animal is placed and cannot escape. The apparatus consists of a vertical cylindrical glass container (height 45 cm, diameter 25cm) filled with tap water at 23±1°C. The water volume is enough to ensure that the animals can not touch the bottom of the container with their hind paws. The forced swim test was conducted over two days. On the first day, rats were introduced to the cylindrical glass tank filled with water for 15 min (not videotaped), towel-dried and returned to their home cage. Twenty-four hours later, the animals were exposed to the same experimental conditions for 5 min, dried and returned to their home cage. Sessions were videotaped and scored using the software ANYMAZE™ (Stoelting Europa, Co., Ireland). All tests were carried out between 8:00 a.m. to 12:00 p.m. The behavior scored in the forced swim test concerns: (1) immobility- floating with the absence of any movement, (2) latency to be immobile- time duration, (3) swimming, (4) climbing. Offspring was tested at the age of 48 days.

#### Elevated plus maze (EPM)

All parts of the apparatus are made of dark polyvinyl plastic. The open and the closed arms of the maze are 50cm above the floor, 50cm long and 10cm wide. Two tests were running simultaneously. The movements of the rats were tracked with digital camera and the individual sessions were analyzed by computer software ANYMAZETM (Stoelting Europe, Ireland). Mild light was provided by a lamp attached above the open arms of the maze. Each session lasted 5 minutes and was started by placement of the rat in the central area facing the open arms of the maze. After each individual trial, the maze was wiped with a mild detergent. The testing was invariably carried out between 8:00 a.m. and 12:00 p.m. The behaviors scored were: (1) distance traveled in the open arms of the elevated plus maze, and (2) time spent in the open arms of the elevated plus maze. The animals were tested at the age of 27 and 47 days.

#### Y-Maze

The Y-Maze Test is widely used to assess exploratory behaviors, learning, and memory function in rodents and short-term memory (38). The apparatus was made from black Plexiglas (50×16×32 cm) in which the arms were symmetrically separated at 120°. No visual cues were placed inside the maze, but different extra-maze cues were visible from all three arms to enable spatial orientation. During the first trial, each animal could freely explore two arms of the maze for 15 min. and its behavior was recorded with a camera. Subsequently, animals were returned to the home cage for one minute and the maze was cleaned with 70% ethanol. The second trial lasted 5 minutes with animal having the access to all three arms. Sessions were videotaped and scored using the software ANYMAZE™ (Stoelting Europa, Co., Ireland). All tests were carried out between 8:00 a.m. to 12:00 p.m. under dim light condition. The animals were tested at the age of 43 days. Spontaneous alternation behavior (the alternation percentage is calculated by dividing the number of alternations by number of possible triads x 100) is considered to reflect spatial working memory, while the total number of arm entries was considered to reflect spontaneous locomotor activity (39,40).

#### Immunohistochemistry assay

Collection of tissue from offspring was executed at different ages of the animals: in juvenile age (PP31) and in adolescent age (PP50). The animals were killed by cervical dislocation. Brains were extracted (not perfused) and post-fixed with 4% paraformaldehyde for 24 h, cryoprotected in 30% sucrose/phosphate-buffered saline solution for up to 1 week, rapid frozen with liquid nitrogen and kept at −80 °C. Before immunohistochemistry assay, brain tissue was sliced in 40 μm sections on a cryostat (Leica). Tissue sections were stored in antifreeze solution at −15°C. The Ca3, Ca4 and DG of the dorsal hippocampus were assessed for the presynaptic marker synaptophysin (Monoclonal Anti-Synaptophysin, Sigma) as described before (20). Tissue was first treated with 0.6 % H_2_O_2_ for 30 minutes at room temperature and then incubated with 5% Normal Goat Serum (NGS) (Lampire Biological Laboratories) in Tris-Buffered Saline Tween at room temperature for 30 min to decrease probability of non-specific antibody binding, followed by overnight incubation at 4°C in mouse anti-synaptophysin (1:200, Sigma Aldrich). Sections were then incubated at room temperature for 2 h in biotinylated goat anti-mouse (1:200, Vector Laboratories). Brain sections were further processed using the Avidin-Biotin Complex (ABC Elite kit; 1:1000; Vector laboratories) and DAB kit (Vector laboratories). Sections were mounted on Starfrost Advanced Adhesive for IHC (Bamed) dried, dehydrated and coverslipped with Permount (Fisher Scientific).

### Quantification

Sections of the dorsal hippocampus were analyzed for optical densities of synaptophysin. Immunoreactivity for all sections was examined under 40x objective using Leica DM4000M. Photomicrographs were taken for three areas within each analyzed hippocampal region, i.e., CA3, CA4 and DG. The software ImageJ64 (Wayne Rasband, NIH, Bethesda MD, USA) was used for assessment of optical densities for all immunoreactive cells. The relative optical density was defined as the difference in optical density (grey level) after calibration between the area of interest and the background, which was an equivalent area adjacent to the area of interest with minimal staining.

### Data analysis

Normality of data was analyzed by Shapiro-Wilks test. Data without normal distribution were analyzed using Kruskal-Wallis test (STATISTICA 10). Analysis of variance (factorial ANOVA) was used to evaluate differences in the individual variables of tests with normally distributed data (STATISTICA 10). Post-hoc comparisons utilized the Tukey HSD test (p≤0.05). The data were expressed as mean ± standard error mean (S.E.M). The changes with values of p≤0.05 were considered statistically significant. All analysis was done in a blind manner.

## RESULTS

### MOTHERS

#### Grimace test

Rat grimace test scores has been used to evaluate spontaneous pain (41). We observed significant main effect of stress on grimace test scores (F (1, 22) = 5.00; p≤0.05). Post-hoc analysis did not reveal any significant differences (Fig 3).

**Figure 3.**
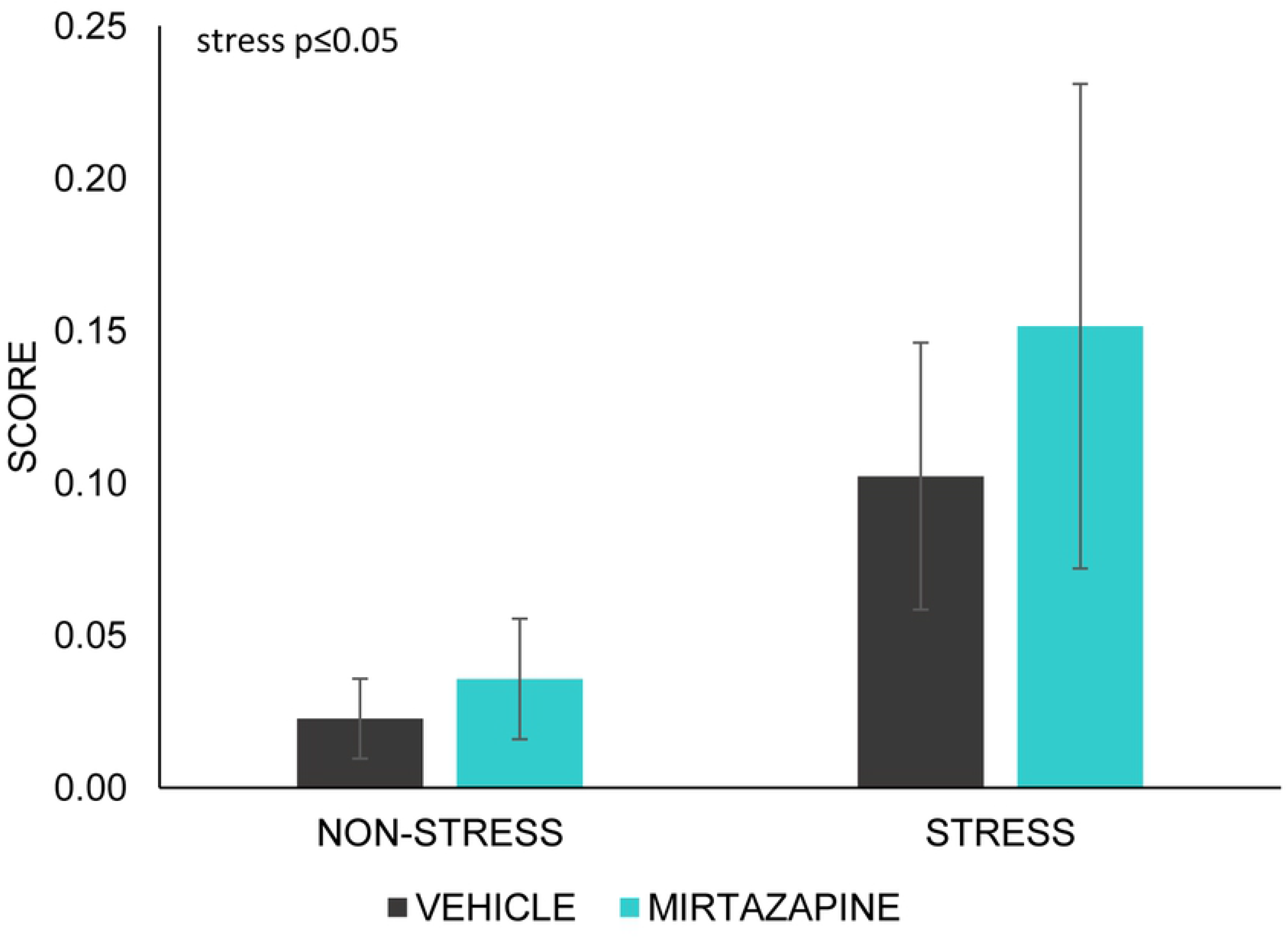
Grimace scale test. Data represent mean ± SEM. n= 6-7 animals/group.

#### Maternal behavior

There was no significant main effect of stress or mirtazapine on total time dams spent nursing (Fig 4A), however, we observed significant main effect of mirtazapine on the percentage of passive nursing and blanket nursing (F (1, 30)= 10.74; p≤0.01). Subsequent post-hoc analysis showed decrease of passive nursing (p≤0.05) and increase of blanket nursing (p≤0.05) in stress × vehicle group as compared to both mirtazapine groups (Fig 4B).

**Figure 4.**
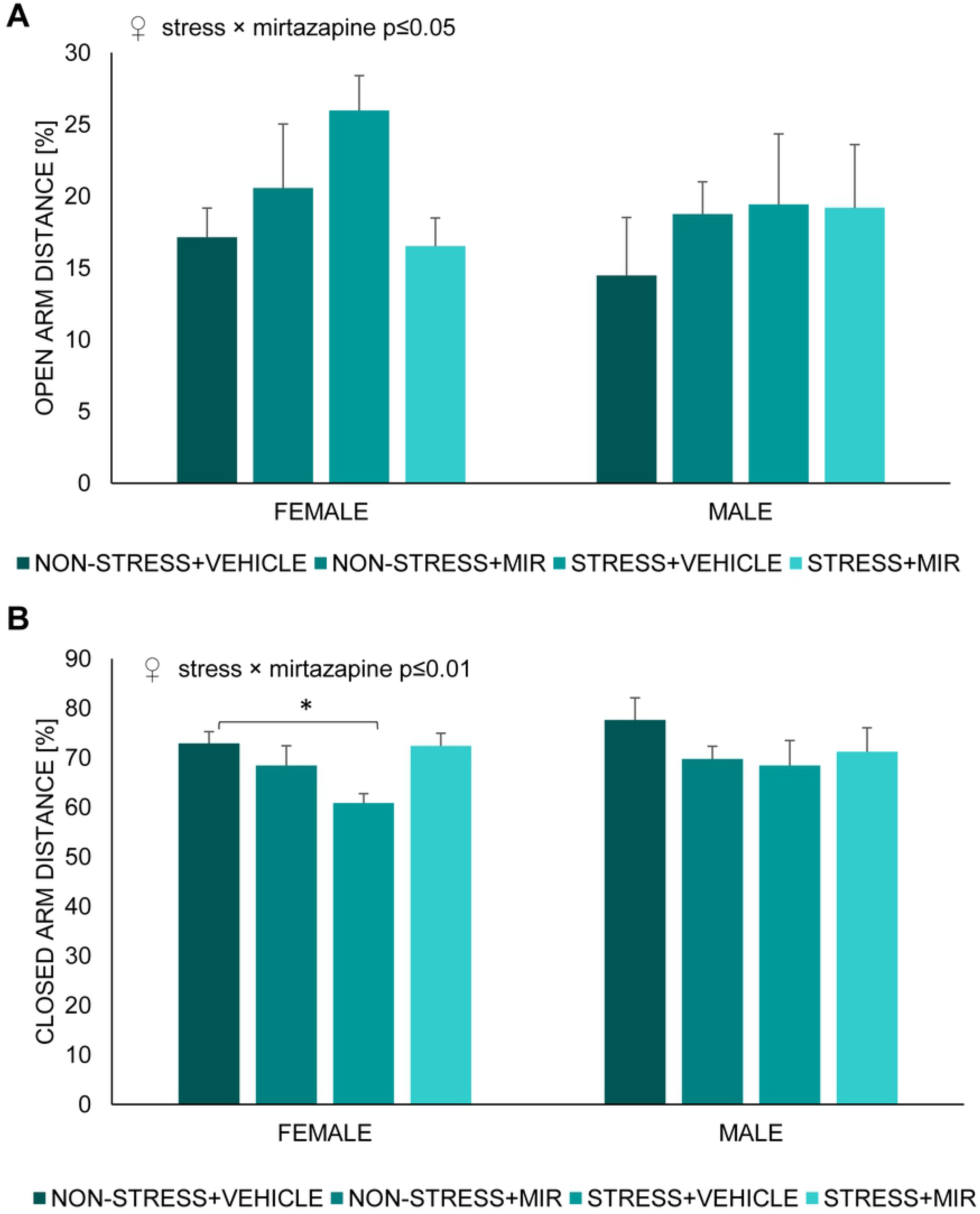
Maternal behavior. (A) total time spent nursing, (B) percentage of passive nursing and blanket nursing out of total time spent nursing. Data represent mean ± SEM. n= 6-10 animals/group. *p≤0.05; m-compared to both mirtazapine groups.

### OFFSPRING

#### Elevated plus maze- juvenile offspring

We did not observe an effect of sex but there was a stress × mirtazapine interaction in females on percentage of distance travelled in the open (F (1, 24)= 4.89; p≤0.05) (Fig 5A) and closed arms (F (1, 24)= 7.82; p≤0.01) (Fig 5B) of EPM, an effect was not present in male offspring. Further analysis showed a decrease in total distance travelled in stress × vehicle females compared to control group.

**Figure 5.**
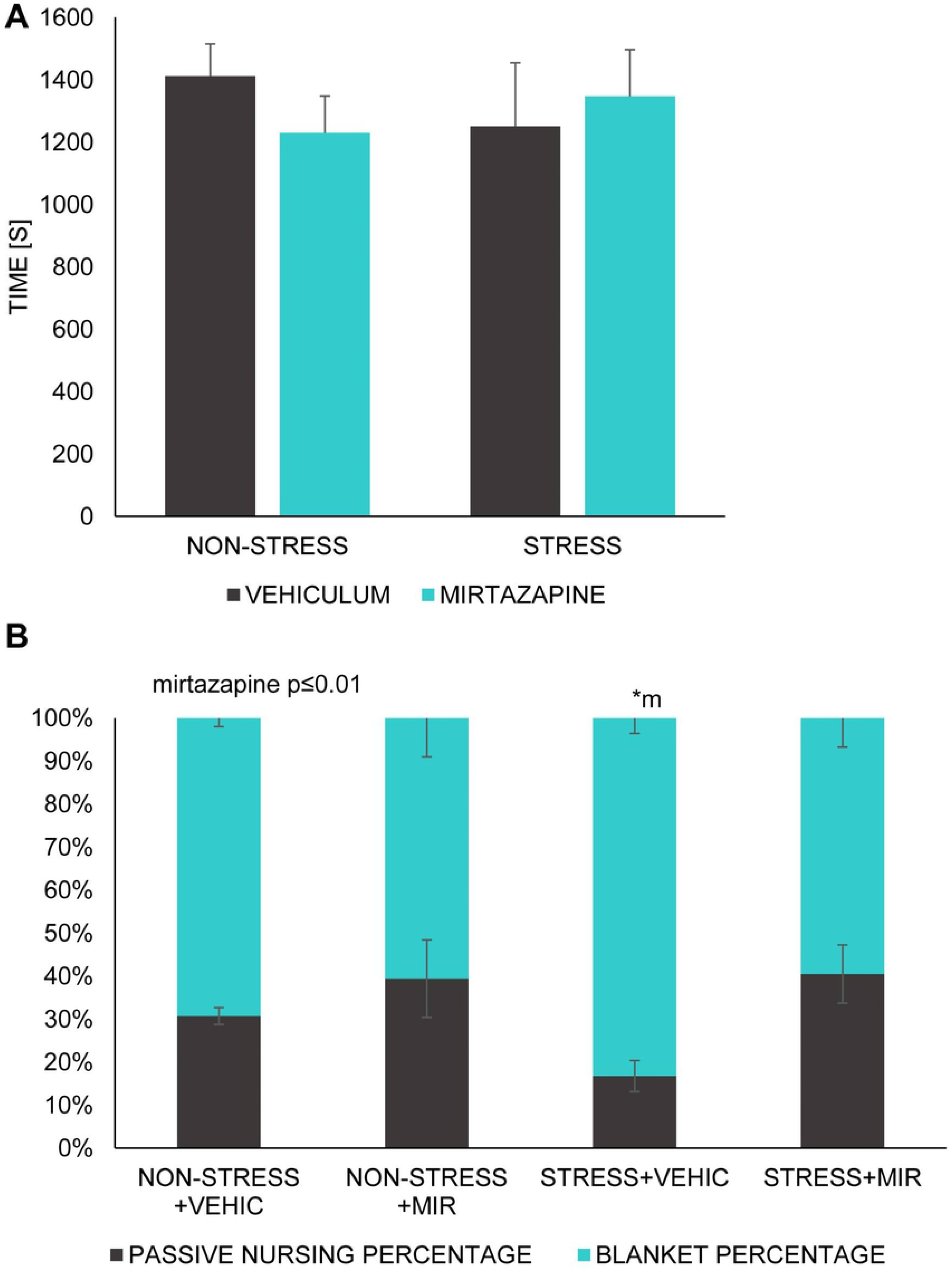
Percentage of distance travelled in individual arms of elevated plus maze by juvenile animals. (A) open arm distance, (B) closed arm distance. Data represent mean ± SEM. n= 7-8 animals/group. MIR-mirtazapine; *p≤0.05.

#### Elevated plus maze- adolescent

Main effect of sex was present in all parameters of elevated plus maze with females being more active (higher total distance travelled) than males (F (1, 62) = 9.04; p≤0.01).

In males, we observed marginally significant main effect of stress × mirtazapine interaction in total distance travelled (F (1, 30) = 3.69; p=0.06). Post-hoc analysis showed significantly decreased total distance travelled in stress × vehicle group (p≤0.05) compared to non-stress × vehicle group. Also, trend for decrease in non-stress × mirtazapine (p=0.07) and stress × mirtazapine (p=0.07) groups compared to non-stress × vehicle group (Fig 6A) was present. In females, we did not observe any differences in total distance travelled (Fig 6A).

**Figure 6.**
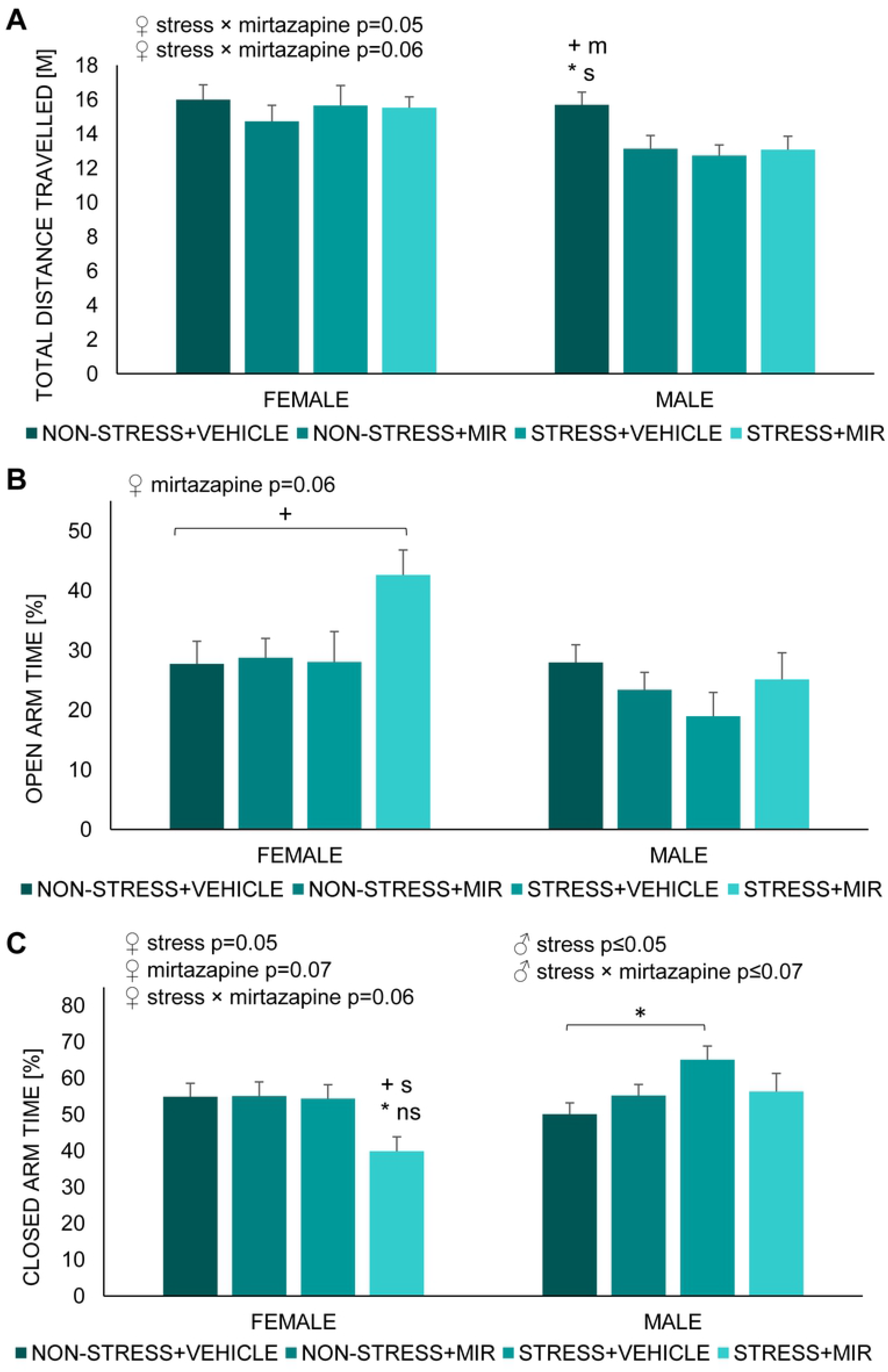
Elevated plus maze of adolescent offspring. (A) total distance travelled, (B) open arm time percentage, (C) closed arm percentage. Data represent mean ± SEM. n= 7-10 animals/group. m-compared to both mirtazapine groups; s-compared to stress × vehicle group; ns-compared to both non-stress groups; + − marginal significance; *p≤0.05.

In males, there were no changes in the percentage of time spent in the open arm (Fig 6B) but we observed a main effect of stress (F (1, 30)= 4.53; p≤0.05) in the percentage of time spent in the closed arm and trend for stress × mirtazapine interaction (1, 30)= 3.37; p=0.07). Post-hoc analysis revealed that animals from stress × vehicle group had significantly increased percentage of time spent in the closed arm compared to non-stress × vehicle group (p≤0.05) (Fig 6C).

In females, we observed marginally significant effect of mirtazapine (F (1, 32)= 3.70; p=0.06) and a trend in effect of stress (F (1, 32)= 3.06; p=0.08) on the percentage of time spent in the open arm. Post-hoc analysis revealed only marginally significant increase in stress × mirtazapine group compared to non-stress × vehicle group (p=0.06) (Fig 6B). There was marginally significant main effect of stress × mirtazapine interaction (F (1, 32) = 3.53; p=0.06) in the percentage of time spent in the closed arm. Post-hoc analysis showed decrease in stress × mirtazapine group significant compared to non-stress × vehicle (p≤0.05) and non-stress × mirtazapine group (p≤0.05) a trend in decrease compared to stress × vehicle group (p=0.08) (Fig 6C).

#### Forced swim test

There were no sex differences in either of the selected forced swim test parameters.

Kruskal-Wallis test showed significant main effect of stress in time spent floating (males: H=7.41; p≤0.01; females: H=6.90; p≤0.01), time spent climbing (males: H=4.47; p≤0.05; females: not significant) and time spent swimming (males: H=6.49; p≤0.01; females: H=10.24; p≤0.001). Further analysis revealed that floating time of stress × mirtazapine males was significantly decreased compared to non-stress × vehicle males (p≤0.01), however floating time of females was not significantly changed between individual groups (Fig 7A). Total swimming time of stress × mirtazapine males was significantly decreased compared to non-stress × vehicle males (p≤0.01) and floating time of stress × mirtazapine females was significantly decreased compared to non-stress × mirtazapine females (p≤0.01) (Fig 7B). Climbing time was not significantly changed between individual groups in either sex (Fig 7C).

**Figure 7.**
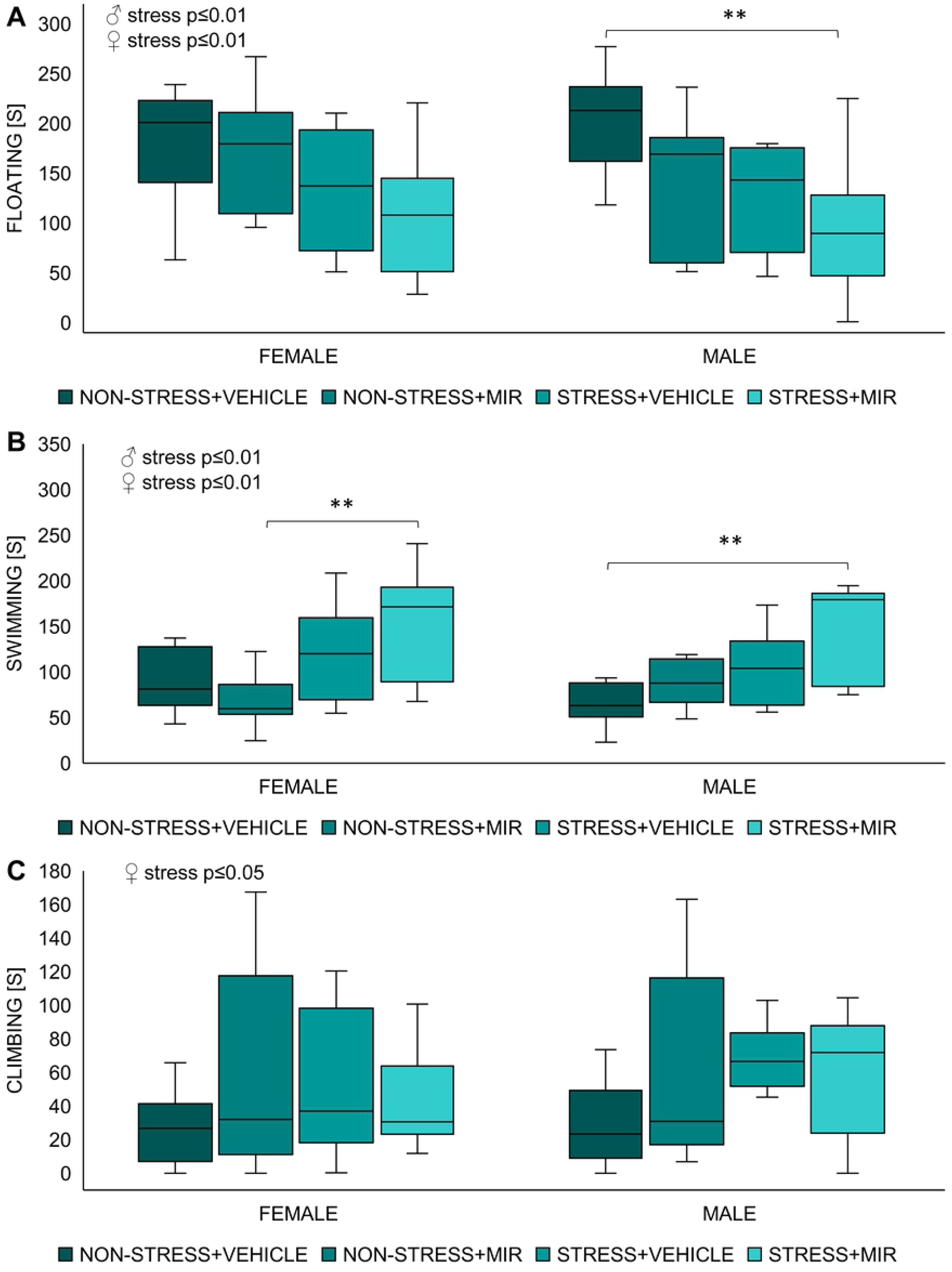
Forced swim test of adolescent offspring. (A) time spent floating, (B) time spent swimming, (C) time spent climbing. n= 7-11 animals/group. **p≤0.01.

#### Y-maze-adolescent

We observed main effect of sex in both motor activity (F (1, 45)= 11.53; p≤0.01) as well as in percentage of spontaneous alterations (F (1, 45)= 4.27; p≤0.05) with females being more active but having decreased percentage of spontaneous alterations compared to males.

In males, a significant main effect of treatment (F (1, 20) = 9.79; p≤0.01) was observed in total distance travelled. Post-hoc analysis revealed marginally significant decrease in non-stress × mirtazapine group compared to non-stress × vehicle group (p=0.07) and significant decrease compared to stress × vehicle group (p≤0.05) (Fig 8A). We did not observe any significant changes percentage of spontaneous alterations (Fig 8B).

**Figure 8.**
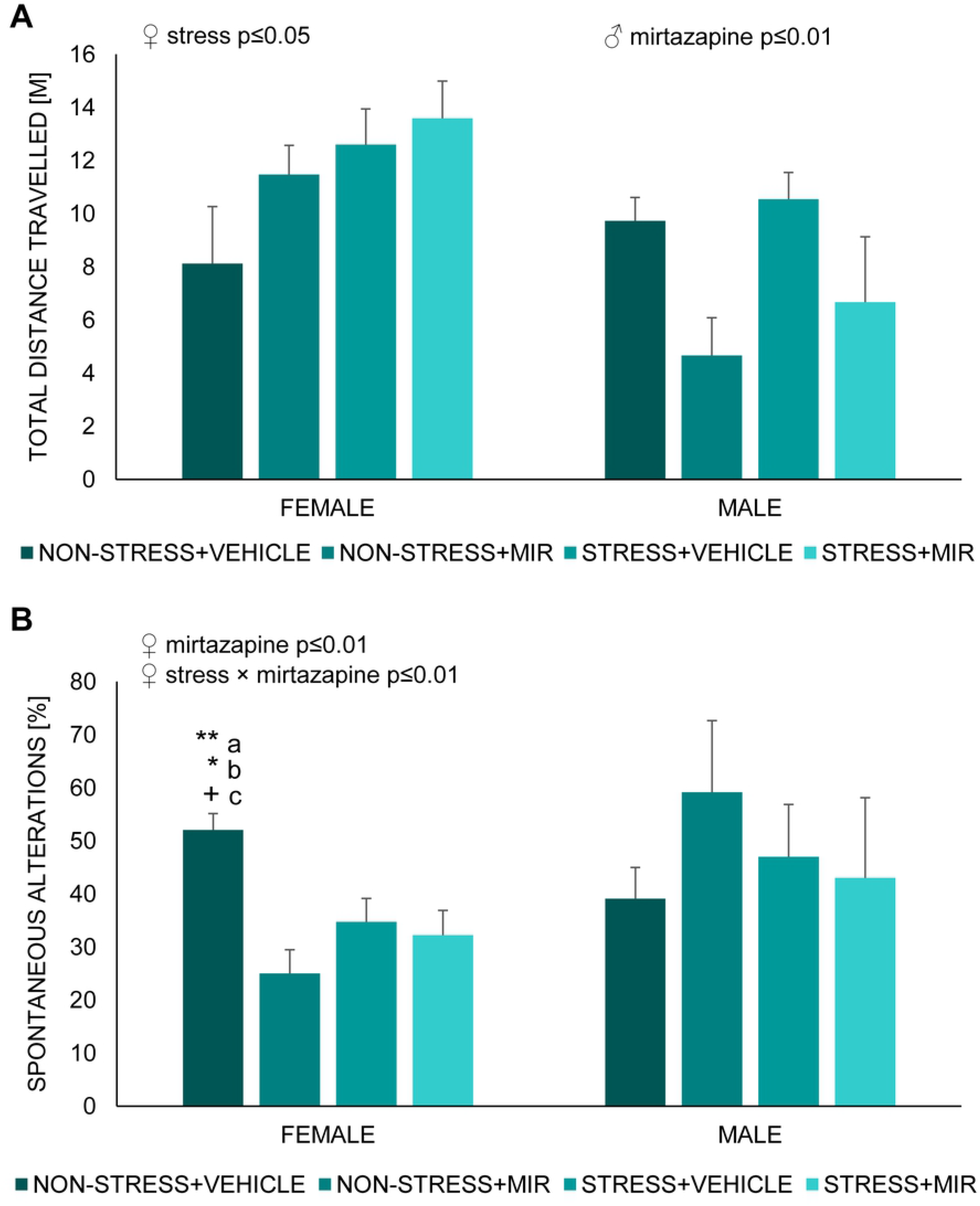
Y-maze of adolescent offspring. (A) total distance travelled, (B) percentage of spontaneous alterations. Data represent mean ± SEM. n= 5-8 animals/group. a-compared to non-stress × mirtazapine, b-compared to stress × vehicle, c-compared to stress × mirtazapine, + − marginal significance, *p≤0.05, **p≤0.01.

In females, a significant main effect of stress was in total distance travelled (F (1, 25) = 4.95; p≤0.05). Post-hoc analysis showed marginally significant increase of horizontal motor activity in stress × mirtazapine group compared to non-stress × vehicle group (p=0.07) (Fig 8A). Further, we observed significant main effect of mirtazapine (F (1, 25) = 11.25; p≤0.01) and mirtazapine × stress interaction (F (1, 25) = 7.81; p≤0.01) in percentage of spontaneous alterations. Post-hoc analysis revealed significant or marginally significant decrease in all groups (non-stress × mirtazapine p≤0.01; stress × vehicle p≤0.06; stress × mirtazapine p≤0.05) compared to non-stress x vehicle group (Fig 8B).

#### Synaptophysin-juvenile

We did not observe any statistically significant changes in the synaptophysin optical density in the hippocampal areas Ca3, Ca4 and dentate gyrus (data not shown).

#### Synaptophysin-adolescent

We found significant main effect of sex present only in area Ca4 of hippocampus (F (1, 47) = 7.23; p≤0.01) (Fig 9A). Further, we observed significant main effect of mirtazapine in hippocampal areas Ca3 (F (1, 19)= 9.01; p≤0.01) (Fig 9B) and Ca4 (F (1, 23)= 5.87; p≤0.05) (Fig 9C) of females. Post-hoc analysis did not reveal significant differences between groups.

**Figure 9.**
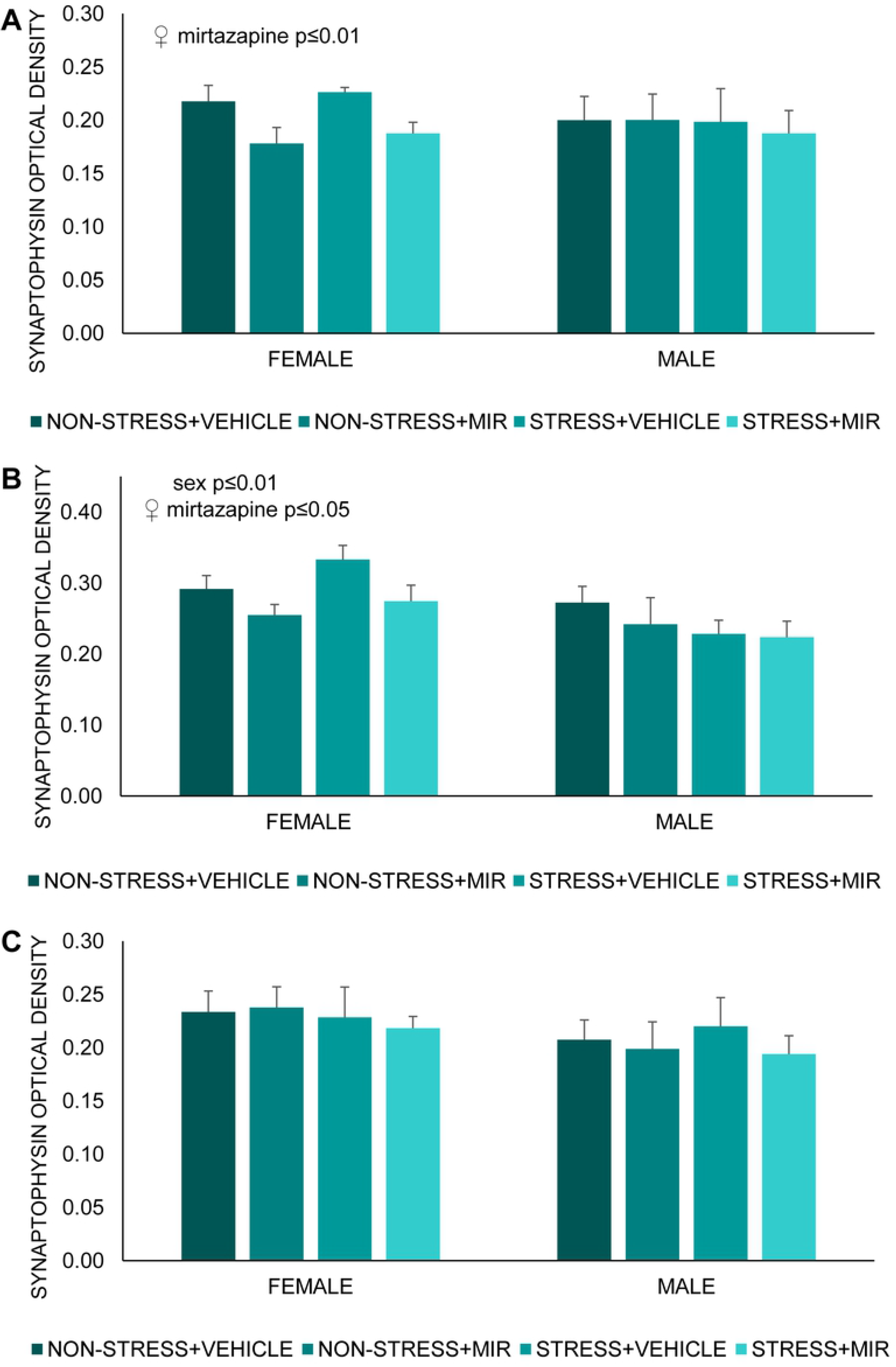
Optical density of synaptophysin in hippocampus. (A) Ca3 area, (B) Ca4 area, (C) dentate gyrus area. Data represent mean ± SEM. n= 5-8.

## DISCUSSION

In the present study, focused on possible implications of pre-gestational chronic stress on the neurodevelopmental and behavioral outputs in the offspring of both sexes during juvenile age and adolescence, we wanted to investigate the consequences of administration of the atypical antidepressant mirtazapine.

We evaluated the spontaneous pain of the mothers by the Rat Grimace Scale (RGS) and we found increased facial expression indicating pain in those mothers, which were exposed to stress schedule. RGS was developed to identify acute and inflammatory pain but it can be applied to a wider range of pain types and chronicity (42,43) and this is supported by our data that suggest ongoing influence of CUS even 5 weeks after the CUS procedure.

Maternal behavior usually emerges close to parturition and after birth, with the female showing a very rapid interest in the new-born (44). Detrimental changes in maternal behavior can generate permanent neurological and social inclusion disturbances in offspring (45). We observed an effect of mirtazapine treatment on the proportion of passive to blanket nursing favoring the latter one, indicating a slight influence of mirtazapine on some aspects of maternal behavior. Mirtazapine acts by antagonizing the adrenergic alpha2-autoreceptors and alpha2-heteroreceptors as well as by blocking 5-HT2 and 5-HT3 receptors. It enhances, therefore, the release of norepinephrine and 5-HT1A-mediated serotonergic transmission (15). Since maternal behavior is strongly regulated by serotonin (46–48), we thus suppose that serotonin activity of MIR can underlie these changes. Shift in maternal behavior in favor of more active blanket nursing caused by antidepressant treatment is in line with our previous results from venlafaxine (serotonin-norepinephrine reuptake inhibitor) (49) as well as results from fluoxetine (selective serotonin reuptake inhibitor) (50).

### Anxiety-like behavior

Offspring of mothers that have experienced stress during gestation may have impaired emotional development due to a dysregulation of the HPA axis, leading to a higher risk of developing a cognitive and/or mood disorders in adulthood (28,51–53). We evaluated the effect of pregestational stress (PS) exposure on intensity of anxiety-like behavior by the elevated plus-maze test, a validated behavioral paradigm based on rodent’s preference of closed to open spaces, with latter presenting a significant anxiety-like behavior inducing condition (54).

In this study, we didn’t find significant sex-dependent differences in anxiety-like behavior of juvenile rats, what was expected due to unmature reproductive system (55). However, female offspring of stressed untreated mothers exhibited decreased anxiety-like behavior that was not present if mothers were treated with MIR. This was changed in adolescence, when females showed decreased anxiety-like behavior if mothers were stressed and treated with MIR. Different results were observed in adolescent male rats, who displayed decreased anxiety-like behavior due to pregestational stress, but mirtazapine treatment of mothers ameliorated this effect. Changes in anxiety-like behavior, due to maternal stress, were previously described by several research groups (28,56). Although there are controversial results regarding effect of chronic pregestational stress on anxiety-like behavior of offspring, possibly due to the different stress paradigm strategies, a great proportion of studies postulate that male offspring whose mothers were submitted to stress before or during pregnancy are more reluctant to enter and spent time in the open arm of the elevated plus maze (57–59). This effect of perinatal mirtazapine treatment on anxiety-like behavior of offspring may be due to the antagonism that mirtazapine exerts on noradrenergic autoreceptors that in turn enhances the noradrenergic transmission opposing the stress-related response (60). However, the research group of Sahoo et al. studied the effects of prenatal exposure to mirtazapine on the postnatal development of rats, with animals spent less time in the open arm, albeit sex was not distinguished (61). Nevertheless, the sex-dependent differences in adaptive behavioral performance to a challenge due to maternal stress may be explained by the influence of individual fetal brain programming, differentially modulating e.g. the responsiveness to a new stimuli, or a hormone arousal during adolescence (62). Recent studies postulate that the sexual hormones, testosterone and estradiol, have a neuroprotective and anti-inflammatory role on cognitive functions (1,63), which is a possible ground for further investigation.

#### Depressive-like behavior

Modified forced swim test, based on the protocol established by Porsolt (64), has been used as a tool to determine the learned helplessness (rodent despair) and therapeutic efficacy of mirtazapine. However, several recent studies postulate that the immobility in this test may not be a sign of a depressive-like state but rather reflection of the individual ability to cope with an acute stressor, explaining the passive phenotype as a result of animal’s difficulties to adapt to external stimuli (65,66).

Our results show an increased active behavior of animals from stressed mothers of both sexes regardless of mirtazapine treatment manifested as less time floating and more time trying to actively escape. Even though several studies declared increased immobility time in animals that have experienced stress during their lives, particularly prenatally, (52,67,68) there are studies showing possibility of prenatal stress inducing adaptive changes contributing to stress resilience later in life (69–71). Active behaviors, such as swimming and climbing, in the rat forced swimming test are differentially regulated by serotonergic and noradrenergic systems (72). Swimming behavior, in this test, is modulated by serotonergic neurotransmission, as Brummelte and colleagues demonstrated a decreased swimming of offspring whose mothers were exposed to high levels of corticosterone while pregnant (73). Also, Vázquez et al. suggested that increased levels of serotonin in control animals after FST may be responsible for shorter time spent immobile and increased time spent swimming, corroborating an effect of the serotonergic system, which in turn modulates dopaminergic projections associated with the reward system and the coping with stress. In this terms, lower levels of serotonin were linked to a stress-induced anhedonic state, while higher levels promote a non-anhedonic state (52). Accordingly, higher levels of serotonin may be a plausible explanation to our results.

#### Spatial memory

We encountered sex-dependent differences in the Y-maze, a behavioral paradigm set to assess the spatial memory performance building on the natural inclination of rodents to explore new environments. Male offspring from mirtazapine treated mothers showed reduced locomotive activity, indicating an influence of treatment on executive capacity. On the other hand, females seemed to be rather affected by the pregestational stress experience of the mothers, which induced a hyperactivity-like response. Such a result may be reflective of hyperactivity or even ADHD described in children of stressed mothers while pregnant (74,75). However, sex differences in reactivity to either pre-gestational stress or perinatal mirtazapine treatment related to potentially hyperactive behavior require further investigation. In children, PS is connected to cognitive, behavioral, and emotional problems such autism and ADHD (76,77). The maternal stress, manifesting as increased cortisol and corticotropin-releasing factor (CRF) levels, affects the fetal development by reprogramming the HPA axis, leading to impaired memory and learning due to the long-lasting effects in the hippocampus (78). Moreover, Deminière et al. had previously described an increased locomotor reactivity to novelty in litters of pre-gestationally stressed mothers, which authors associated with a modified dopaminergic activity in the prefrontal cortex and nucleus accumbens (79). Experience-dependent plasticity of hippocampal neurons and adult neurogenesis, processes of importance for optimal learning or memory formation and processing, are modulated through epigenetic mechanisms, which induce long-lasting changes determining the ability of individuals to cope with adverse situations (80,81).

In our study, adolescent females had impaired spatial memory, represented by decreased percentage of spontaneous alternations, as a result of maternal stress as well as mirtazapine treatment. However, this effect was not present in males. Conrad et al. studied the effects of chronic stress on spatial memory performance of adult rats concluding that stress impaired the spatial learning and memory, hippocampi-dependent spatial tasks. Authors also resolved that females seem to be less vulnerable to these hippocampi-dependent memory deficits (38,82). Nevertheless, they didn’t study the effects in the offspring. Otherwise, female rats have been linked to more active response in novel environments (30,83,84) and, in line with our results, pregestational stress as well antidepressant treatment highlighted this response with effect of mirtazapine being more pronounced in animals without pregestational stress exposure. Similar results have been postulated in association with other antidepressant such as fluoxetine, bupropion or citalopram (74,85). Nevertheless, our results show a clear influence of pre-gestational stress and antidepressant treatment in the onset of behavioral tasks, which may be driven by the differential strategies used by each sex to cope with adversity. Moreover, we could take into account that we evaluated animals during adolescence, which is known as a time of increased physiological and psychological changes, which make them more vulnerable and unpredictable to external influences (86–88).

#### Synaptophysin analysis

Synaptophysin is used as a marker of synaptic plasticity and synaptic nerve terminal density (89). We analyzed optical density (OD) of synaptophysin, a calcium-binding glycoprotein widely distributed in the presynaptic vesicle membrane, that is required for vesicle fusion and neurotransmitter release, in the CA3, CA4 and DG areas of the hippocampus. Results of synaptophysin density in our study were age-dependent with no changes in juvenile offspring. However, with onset of adolescence we observed prominent effect of sex with synaptophysin OD males remaining intact under all conditions while females being influenced by maternal mirtazapine treatment. This effect was regionally specific as changes were observed only in CA3 and CA4 areas but not in DG. Sex differences can be seen in the neural plasticity at DG-CA3 synapses. In short, mossy fibers evoke larger population spikes in CA3 pyramidal neurons in females during proestrus and estrus relative to males, while mossy fibers in males have stronger synaptic connections to CA3 neurons than females (90). Limitation of our study is that we didn’t evaluate estrus cycle phase before extraction of the brains. There is very limited research available on maternal antidepressant treatment in association with synaptophysin OD. Fluoxetine, a SSRI, has been proven to impact synaptophysin OD of CA3 in the same sex- and age-dependent manner as seen in our study (31). Previous studies have encountered a reduced expression of synaptophysin in the hippocampus as whole or in DG, no research is done on other specific regions, at different ages (PP7, PP14, adult) of pre-gestationally stressed pups (91–93), suggesting the involvement of early-life stress in modulation of synaptic plasticity, which could lead to development of neuronal deficits during adulthood (94,95). Previous studies suggested a regulatory role of ovarian hormones in the developing brain (94) explaining the differential effect of stress on limbic synaptic plasticity in female rats. However, there are differences between our study compared to studies previously mentioned. Our model comprises of CUS prior to gestation and we tested the offspring in adolescent age, when the reproductive system of the offspring is not fully developed, so we can’t rule out other stress related changes in later life that were not seen in this study.

## CONCLUSIONS

Neurobiology of depression remains mostly unexplained with several theories trying to identify its etiology including the impairment of monoaminergic systems coupled with an altered neural plasticity, dysregulation of the HPA axis or the immunological response. Maternal depression as well as antidepressant treatment thus may lead to a broad spectrum of neurodevelopmental changes in the offspring. Our results suggest mirtazapine induced alterations in maternal behavior and several sex- and age-dependent changes in neurobehavioral development of offspring caused by either or combination of prenatal mirtazapine treatment and chronic pre-gestational stress.

## ACKNOWLEDGMENTS

The study was supported by the VEGA Scientific Agency (VEGA 2/0124/19) and APVV grant of Slovak Republic (APVV-19-0435). The authors confirm that there is no conflict of interest.

